# Isolation of Neural Stem Cells Using Platelet Derived Growth Factor C

**DOI:** 10.1101/2025.04.29.650770

**Authors:** Hiba K. Omairi, Mathieu Meode, Sarah Sait, Kyle M. Heemskerk, Sophie Black, Jennifer A. Chan, J. Gregory Cairncross

## Abstract

Here, we show that Platelet-Derived Growth Factor C (PDGFC) supports the isolation of neural stem cells (NSCs) from the murine subventricular zone (SVZ) and maintains them in quiescent and slowly proliferating states. We also show that NSCs isolated using PDGFC can be induced to proliferate rapidly when switched to media supplemented with Epidermal Growth Factor and Fibroblast Growth Factor (EGF/FGF), the gold standard growth condition for NSCs. Although patterns of gene expression in NSCs isolated in PDGFC or in EGF/FGF are similar, a comparative analysis of quiescence genes reveals that NSCs in PDGFC are more like SVZ tissue than are NSCs maintained in EGF/FGF. In addition, NSCs isolated using PDGFC transition to oligodendrocyte progenitor cells (OPCs) when FGF is added to PDGFC and have an expression profile indistinguishable from OPCs that are isolated traditionally using PDGFA/FGF.

## Introduction

Reynolds and Weiss reported that cells from the adult mouse striatum cultured in EGF retain the capacity to divide and differentiate into neurons and astrocytes (Reynolds and Weiss, 1992). This finding demonstrated that stem-like cells persisted in the postnatal brain and raised the intriguing possibility that they might be activated to repair the brain after injury. Thirty years later, clinically significant brain repair has yet to materialize, but the finding of stem-like cells in the adult mammalian brain has assumed renewed significance in the context of cancers of the central nervous system. It is now widely held that the deadly brain cancer of adults, Glioblastoma Multiforme (GBM), arises from stem or progenitor cells that persist in the mature brain (Liu et al., 2011). Indeed, some have speculated that cancer-predisposing alterations residing in quiescent stem cells constitute the first step in the pathogenesis of GBM (Lee et al., 2018).

Consistent with this hypothesis, we reported that SVZ-derived murine cells that are P53 null (or heterozygous for P53) become exogenous growth factor independent and tumorigenic during chronic exposure to Platelet-Derived Growth Factor A (PDGFA) while acquiring copy number alterations that closely resemble those seen in human GBM cells and tumors (Omairi et al., 2023). In contrast to null cells, we observed that P53 wild type SVZ cells neither divided nor survived in PDGFA (Bohm et al., 2020). These different P53-associated behaviors of SVZ cells prompted us to explore how they might respond to PDGFC. We chose PDGFC, because like PDGFA, PDGFC exerts its biological effects through PDGFRα (Fredriksson et al., 2004). Rather than dichotomous behaviors – transformation *vs*. cell death – both null and wild type SVZ cells formed small spheres in PDGFC. We then examined the composition of these spheres focusing on wild type cells.

## Results

### Murine multipotent NSCs can be isolated from the SVZ using PDGFC

SVZ-derived cells cultured in defined media supplemented with PDGFC only had a distinctive phenotype; they gradually formed small slowly enlarging spheres. Their behavior in PDGFC contrasted with that observed in EGF/FGF where, as expected, they proliferated rapidly and continuously and formed large spheres (Bohm et al., 2020; Omairi et al., 2023). To explore the potential physiological relevance of primary SVZ cell cultures in PDGFC, we examined the expression of *Pdgfc* in the SVZ of young adult mice relative to other brain regions by RNA sequencing. Analysis of fresh tissue revealed that *Pdgfc* was expressed in the SVZ and that levels of expression were higher than in the cerebral cortex or subcortical white matter (Fig. 1A). This result suggests that stem and progenitor cells residing in the SVZ are exposed to PDGFC.

**Figure 1.**
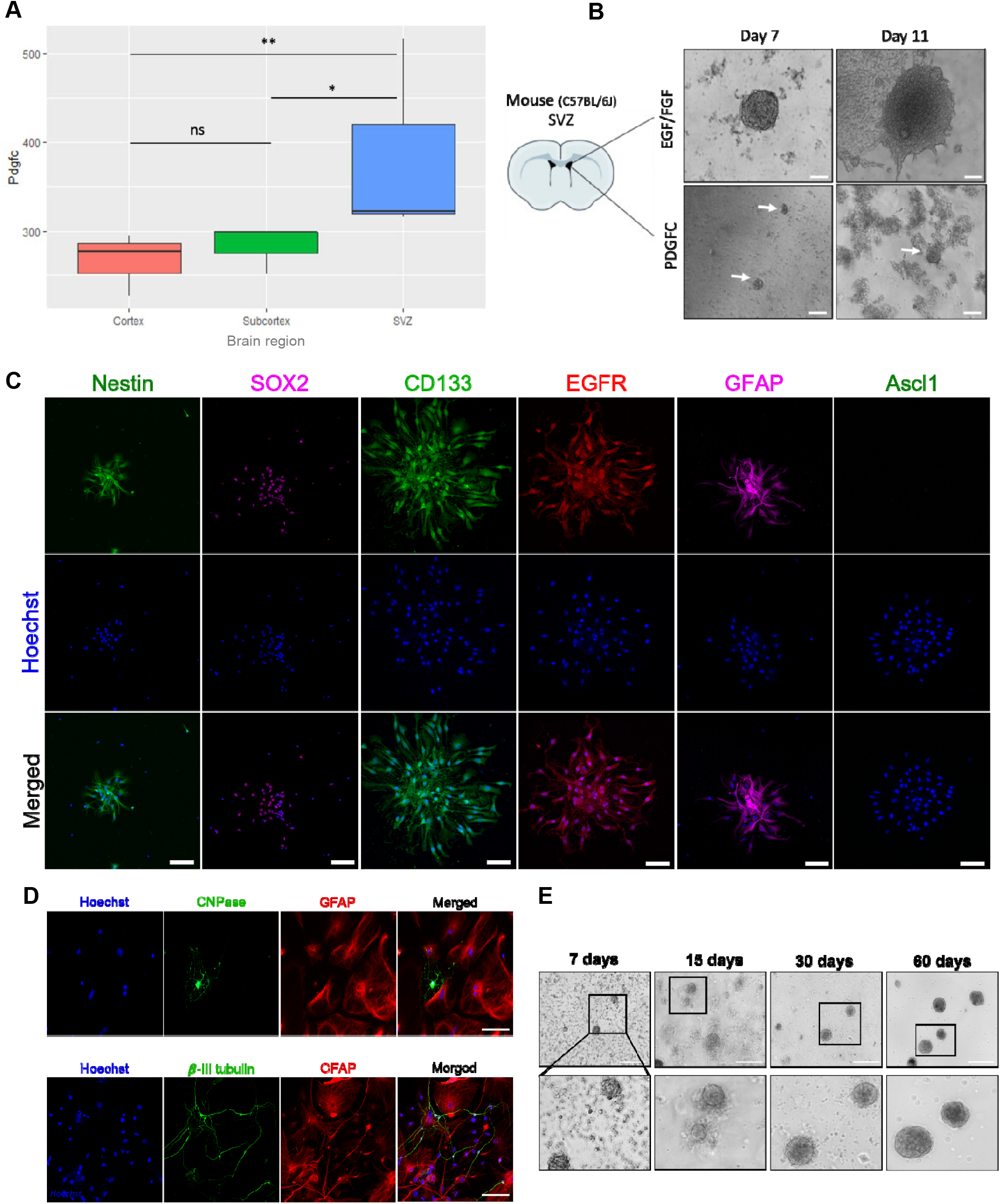
SVZ-derived neural stem cells isolated using PDGFC. **(A)** Boxplot showing PDGFC expression in different regions of the mouse brain: cortex, subcortex and SVZ (n=3) (ns = p > 0.05; * =p≤0.05; **=p≤0.001). **(B)** Representative brightfield images of sphere-forming cultures established in EGF/FGF and PDGFC after 7 and 11 days. Arrows point to small spheres in PDGFC. **(C)** Representative immunofluorescence images of SVZ-derived cells in PDGFC stained for the stem cell markers Nestin, SOX2, CD133, EGFR and GFAP and the Type C cell marker Ascl1 [Mash1] [Scale bar: 50 μm]. **(D)** Representative brighfield images showing spheres isolated in PDGFC at 7, 15, 30 and 60 days after culture initiation [Scale bars :100 µm]. **(E)** Representative immunofluorescence images of cells isolated in PDGFC, differentiated in 1% FBS for 6 days, and stained for markers of astrocytic (GFAP), oligodendroglial (CNPase) and neuronal (Beta-III-tubulin) differentiation [Scale bar: 100 μm].

We then compared primary SVZ cultures isolated in PDGFC or EGF/FGF. Within one week, spheres were visible in both growth factor conditions (Fig. 1B), but those that formed in PDGFC were smaller (53.1±10.77mm *vs*. 99.05±23.68mm; Fig. S1A). Four days later, spheres in PDGFC had enlarged slightly whereas those in EGF/FGF were significantly larger (67.45±15.78mm *vs*. 195.33±46.19mm; Fig. S1A), and soon thereafter developed necrotic centers and required passaging. Based on these observations, we concluded that PDGFC, like EGF/FGF, supports sphere formation in SVZ-derived cells, but not the rapid proliferation of cells that is characteristic of the gold standard growth condition for neural stem cells, EGF/FGF. Furthermore, spheres isolated from the SVZ using PDGFC remained viable for up to 100 days and capable of rapid continuous expansion in EGF/FGF. To our knowledge, no member of the PDGF family of ligands has been shown to support the formation of spheres *in vitro*.

To investigate the nature of these sphere-forming cells and compare them to cells that populate the rapidly enlarging spheres in EGF/FGF, we analyzed their marker profiles using immunofluorescence. All cells in both sets of spheres were positive for the neural stem cell markers SOX2 (i.e., Sex-determining region Y-box 2) and Nestin (Fig. 1C, Fig S2A). Cells in the PDGFC spheres also expressed markers associated with type B cells of the SVZ, namely CD133, EGFR and GFAP, but did not express Mash1 (Ascl) a marker of more differentiated type C cells (Fig. 1C) (Doetsch et al., 1999). To test whether the sphere-forming cells in PDGFC had the capacity for multilineage differentiation, a defining characteristic of stem cells, we induced differentiation using 1% FBS and assessed their morphology and expression of lineage associated markers. Cells in the PDGFC spheres assumed an adherent and stellate morphology and expressed increasing levels of β-III-tubulin, CNPase and GFAP, markers of neuronal, oligodendroglial and astrocytic differentiation, respectively (Fig. 1D, Fig S2B). Sphere-forming cells in EGF/FGF responded similarly. These findings led us to conclude that SVZ-derived cells that survive and form spheres in PDGFC display the characteristics of multipotent NSCs.

To further study the behavior of SVZ cells in PDGFC, we measured changes in sphere size over 2 months (Fig. 1E, S1B). In PDGFC, sphere diameter fluctuated within narrow limits rarely exceeding 100nm. Spheres in PDGFC did not develop necrotic cores or require passaging, leading us to assess the rate of proliferation of cells in PDGFC *vs*. EGF/FGF spheres.

### PDGFC isolated cultures contain slowly proliferating and quiescent stem cells

First, we analyzed the cell cycle profile of NSCs isolated in PDGFC *vs*. EGF/FGF over 48hrs and found significantly more cells in G1 in the PDGFC condition and significantly fewer in S-phase (Fig. 2A). We then measured EdU incorporation over 24hrs in 25-day-old spheres in PDGFC and observed that few cells were positive for EdU (Fig 2B), whereas EdU incorporation increased dramatically and significantly when spheres in PDGFC were switched to EGF/FGF for 5 days (Fig 2B). This result led us to ask whether the non-proliferating cells in PDGFC had exited the cell cycle. To test this possibility, we co-stained age-matched PDGFC and EGF/FGF spheres for expression of the G0 marker, P27 (KIP1), and the proliferation marker, Ki67 (Fig 2C, E). We found that spheres in PDGFC contained a significantly higher number of P27 positive cells and a significantly lower number of Ki67 positive cells than spheres in EGF/FGF (44.8% *vs*. 0.5% and 83.4% *vs*. 22% respectively). Thirty-three percent of cells in PDGFC and sixteen percent of cells in EGF/FGF were negative for both P27 and Ki67 (Fig 2C).

**Figure 2.**
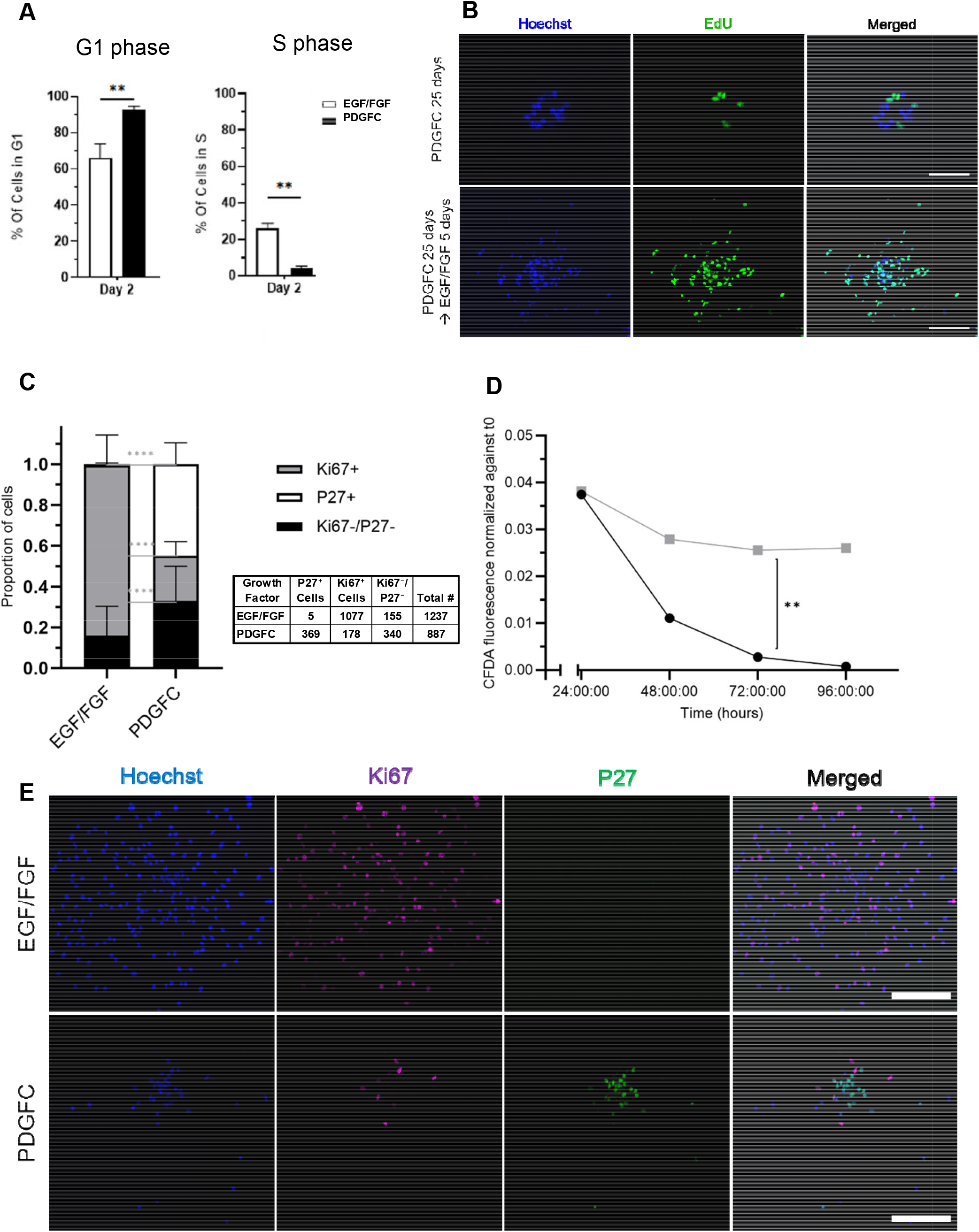
PDGFC cultures contain slowly proliferating and quiescent stem cells. **(A)** Propidium iodide (PI) staining and cell cycle analysis of NSCs cultured in EGF/FGF (control) or PDGFC for 48 hours (n =3, **=p ≤0.01). **(B)** Representative immunofluorescence images of EdU labelled NSCs in PDGFC (day 25) and after switching to EGF/FGF for 5 days (25+5 days). **(C)** Bar graph showing the proportion of P27-positive, Ki-67 positive and double negative cells in EGF/FGF and PDGFC cultures: 1237 cells were counted in the EGF/FGF condition and 887 in PDGFC (n=3; ****=p≤0.0001). **(D)** Flow cytometry analysis using CFDA-SE cell tracking stain on NSCs cultured in PDGFC or EGF/FGF over 4 days. CFDA fluorescence intensity was measured after 24, 48, 72 and 96 hours in both growth conditions. The intensity of CFDA indicates the rate of proliferation as reflected by stain partitioning during cell division (n=5, **=p≤0.01). **(E)** Representative immunofluorescence images of NSCs in PDGFC and EGF/FGF (control) stained for the proliferation marker, Ki67 (Magenta) and the G0 marker, P27 (Green) 7 days after culture initiation [Scale bars: 100 μm].

To further assess cell division in the PDGFC and EGF/FGF conditions, we tracked cell proliferation using the permeable stain, carboxyfluorescein diacetate succinimidyl ester (CFDA-SE). After exposure to CFDA-SE, we evaluated cell fluorescence intensity in cells in PDGFC *vs*. EGF/FGF after 24, 48, 72 and 96hrs (Fig 2D). CFDA-SE was undetectable after 4 days in the EGF/FGF growth condition consistent with a high rate of cell division. However, in PDGFC, a reduction in fluorescence after 2 days was followed by stable fluorescence intensity thereafter suggesting that cells in spheres in the PDGFC condition had a lower rate of proliferation (Fig. 2D). These results also suggest that spheres isolated in PDGFC contain two populations of cells, a minority population that is proliferating and a larger population that is non-proliferative but can be induced to enter the cell cycle (i.e., exit G0) when exposed to EGF/FGF.

### Neural cells of interest can be derived from NSCs in PDGFC

Having observed that multipotent NSCs can be cultured directly from the SVZ using PDGFC (Fig. 1), we inquired whether other populations of neural cells could be derived from this stem cell pool. After preparing 30-day-old spheres in PDGFC, we switched them to EGF/FGF to observe their rate of growth; to PDGFA to assess their behavior in a growth factor that also activates PDGFRα; and to PDGFC+FGF to test whether this growth condition would mimic the combination of PDGFA+FGF that is widely used to isolate and propagate OPCs (Fig S3A) (Chen et al., 2007). One week later, spheres switched to EGF/FGF were significantly larger and required passaging, those switched to PDGFA had deteriorated and were non-viable after 20 days, and those switched to PDGFC+FGF contained enlarging spheres (Fig. S3A). Rapidly proliferating spheres in EGF/FGF reverted to a quiescent phenotype when returned to media supplemented only with PDGFC (Fig. S3B).

Since PDGFC and PDGFA activate the same receptor and quiescent spheres in PDGFC begin to proliferate rapidly when FGF is added to PDGFC, we asked whether PDGFC could be used in combination with FGF to isolate OPCs directly from the SVZ. To that end, we dissected the SVZ from young adult mice and established primary cultures in media supplemented with PDGFC+FGF (experimental condition) or PDGFA+FGF (gold standard condition). ^10^ Within 7 days, large SOX2 and Nestin-positive spheres were visible in both conditions (Fig. S4A). In addition, spheres in PDGFC+FGF were positive for the OPC markers PDGFRα and NG2 and were indistinguishable from spheres in PDGFA+FGF (Fig. S4A, B). These data suggest that PDGFC can substitute for PDGFA when isolating OPCs from the SVZ.

### NSCs can be isolated from the cerebral cortex using PDGFC

We also observed that sphere-forming cells could be isolated from the cerebral cortex of young adult mice using PDGFC (Fig. S3C). Lower yields (fewer spheres per volume of fresh tissue harvested) were obtained from the cerebral cortex than from the SVZ, but like spheres derived from SVZ cells, those from the cerebral cortex were positive for SOX2 and Nestin (Fig, S3D), assumed an adherent stellate morphology in FBS and expanded in EGF/FGF.

### Transcriptomic analysis of NSCs and OPCs isolated using PDGFC

To further examine these neural cell cultures, we performed bulk-RNA sequencing of one-week old NSCs isolated in PDGFC or EGF/FGF. Data were compared to freshly dissected SVZ tissue (Day 0). In PDGFC *vs*. EGF/FGF, 5,741 genes were differentially expressed (FDR <0.05; Data S1). Compared to SVZ tissue, 8,146 genes were differentially expressed in PDGFC and 11,131 in EGF/FGF. Unsupervised clustering and principal component analysis (PCA) showed that PDGFC-isolated NSC populations clustered separately from those isolated in EGF/FGF (Fig. 3A, B). These analyses also revealed that SVZ tissue samples clustered together, remote from the *in vitro* samples, but closer to the samples from the PDGFC condition (Fig. 3A, B). To obtain a better understanding of similarities between the EGF/FGF, PDGFC and SVZ conditions, we generated a sample-to-sample distance heatmap (Fig. 3C). These data show that the *in vitro* PDGFC growth condition was closer to the SVZ tissue of origin than was the EGF/FGF condition. With respect to OPC cultures, all samples (isolated from the same mouse) clustered tightly together whether cultured in PDGFC+FGF or PDGFA+FGF (Fig. 3A).

**Figure 3.**
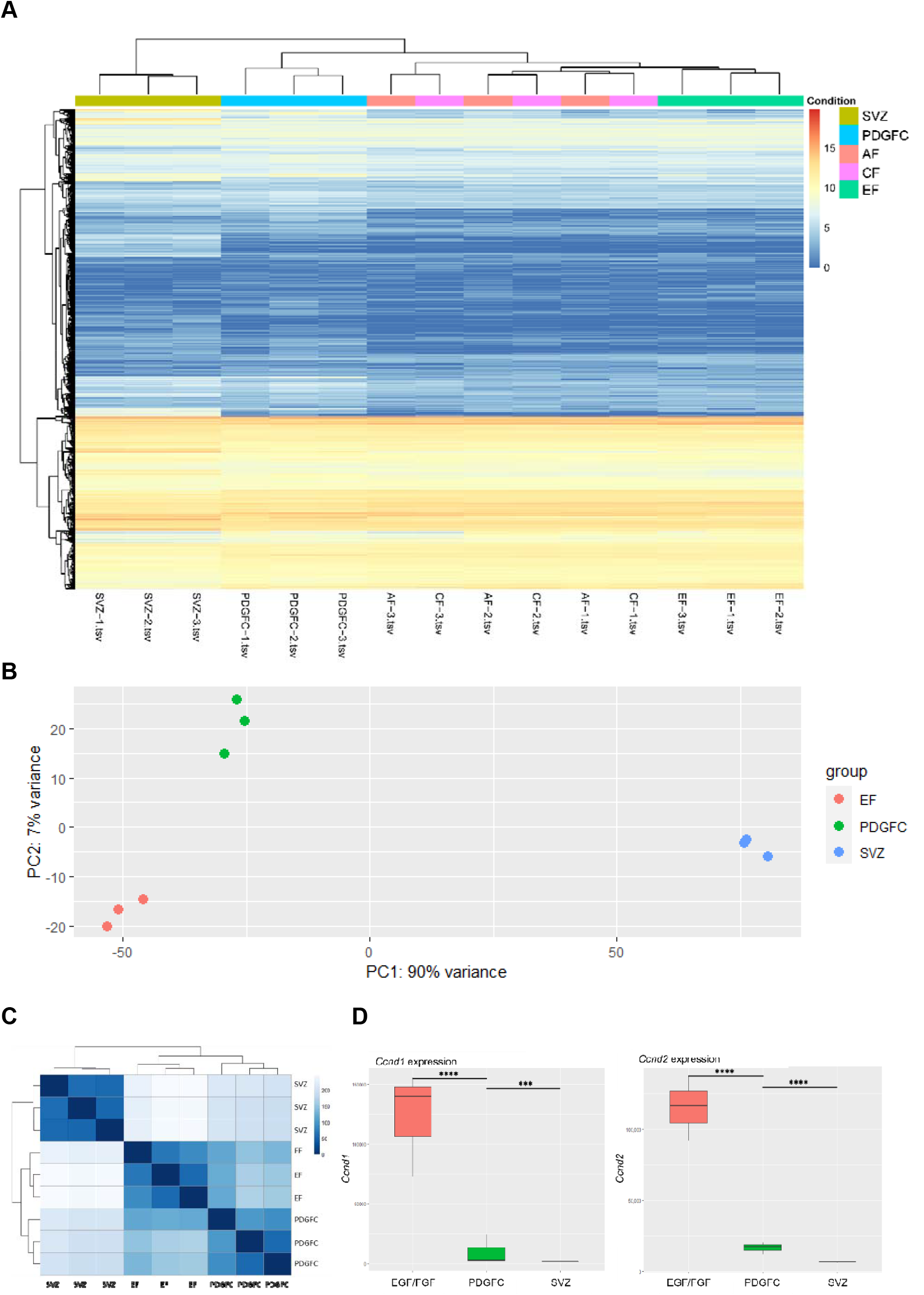
Transcriptomic analysis of SVZ-derived cells isolated in PDGFC. **(A**) Heatmap showing total gene expression and unsupervised sample clustering in fresh *ex vivo* SVZ tissue and SVZ-derived cultures isolated in EGF/FGF, PDGFC, PDGFC+FGF and PDGFA+FGF for 7 days (n=3 for each condition). **(B)** PCA plot containing fresh SVZ tissue and SVZ-derived cultures isolated in EGF/FGF or PDGFC (n=3). **(C)** Sample-to-sample distances heatmap between fresh SVZ tissue and SVZ-derived cultures isolated in EGF/FGF or PDGFC (n=3). **(D)** Boxplots showing the expression of Cyclin D1 and D2 (Ccnd1 and Ccnd2, respectively) in cells isolated using EGF/FGF or PDGFC for 7 days and compared to fresh SVZ tissue expression (n=3; ***=p≤0.001; ****=p≤0.0001).

The different proliferative behaviors of NSCs in PDGFC and EGF/FGF led us to study the expression of two important cell cycle regulators, cyclin D1 and cyclin D2 (*Ccnd1/2*) in both growth factor conditions and in SVZ tissue. We found that *Ccnd1/2* were significantly overexpressed in the EGF/FGF condition *vs*. PDGFC (p-value<0.0001; Fig. 3D). Similarly, the proliferation-promoting immediate early response genes *C-Fos* and *C-Jun* were significantly overexpressed in EGF/FGF *vs*. PDGFC (Fig. 4A), a finding that is consistent with higher overall activation of the G1/S transition in EGF/FGF than PDGFC, and with our Ki67 data (Fig. 2C, E). As expected, *Ccnd1, Ccnd2, C-Fos* and *C-Jun* were significantly overexpressed in both the EGF/FGF and PDGFC growth factor conditions than in SVZ tissue (Fig. 3D, 4A).

**Figure 4.**
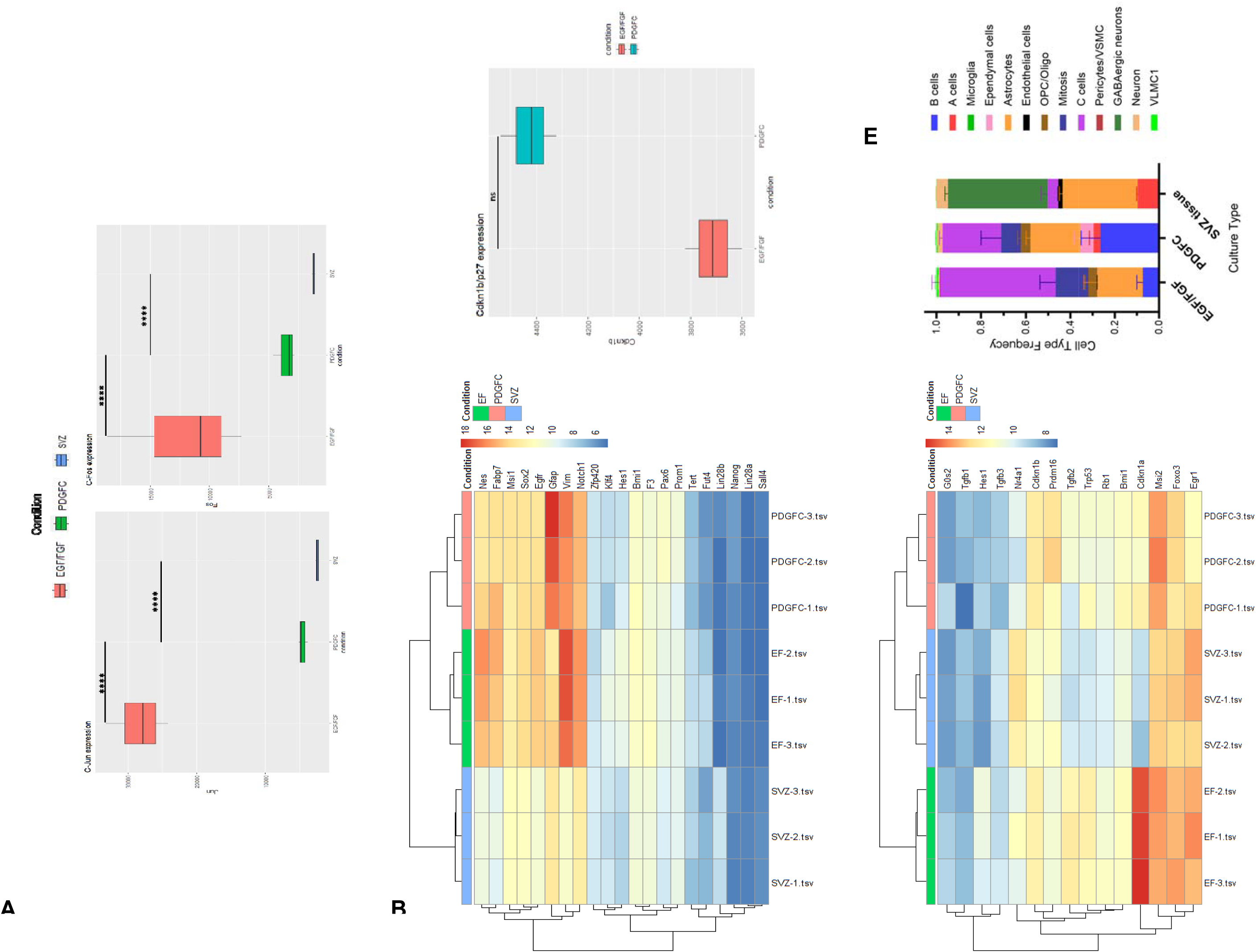
Transcriptomic analysis of differentially expressed cell cycle, stemness and quiescence genes in SVZ-derived cultures. **(A)** Boxplots showing the expression of immediate early response genes c-Jun and c-Fos in EGF/FGF, PDGFC and SVZ tissue (n=3, ****=p-value≤0.0001). **(B)** Heatmap showing expression of 21 genes associated with stemness in EGF/FGF, PDGFC and SVZ tissue (values in normalized and log-transformed counts, vsd). **(C)** Boxplot showing Cdkn1b/p27 expression in EGF/FGF and PDGFC (n=3, ns: not significant). (**D**) Heatmap showing expression of 15 genes associated with quiescence in EGF/FGF, PDGFC and SVZ tissue (values in normalized and log-transformed counts, vsd). **(E)** Cell type composition of SVZ tissue and cultures in EGF/FGF or PDGFC (n=3).

Furthermore, we compared stemness and quiescence gene signatures in the EGF/FGF, PDGFC and SVZ samples. Using a set of 21 genes associated with stemness, NSCs isolated in EGF/FGF and those isolated in PDGFC clustered together (Fig. 4B). However, when considering 15 genes associated with quiescence, NSCs in PDGFC and SVZ tissue were grouped more closely (Fig. 4D). As seen in figures 2C and 2E, the expression of *Cdkn1b* (coding for P27) was higher in the PDGFC condition than in EGF/FGF, a difference that was not statistically significant (Fig. 4C).

Lastly, we used our bulk RNA-Seq data and published single-cell RNA-Seq data from the adult mouse SVZ (Cebrian-Silla et al., 2021) to perform a deconvolution analysis (figure 4E). Quiescent NSCs called type B cells were significantly enriched in the PDGFC condition compared to the SVZ tissue or cells in EGF/FGF (p-values <0.0001 and =0.0004, respectively). We also confirmed a higher proportion of cells in mitosis in EGF/FGF *vs*. SVZ tissue (p-value = 0.0121), a difference that was not significant in PDGFC *vs*. SVZ (p-value = 0.2982).

## Discussion

As we continued to explore of effects of the PDGF ligands on cells derived from the sub-ventricular zone of young adult mice (Bohm et al., 2020; Omairi et al., 2023), we observed that PDGFC supports the isolation of neural stem cells with a distinctive behaviour. In defined media supplemented only with PDGFC, sphere-forming, SOX2+/Nestin+, multipotent neural cells were successfully isolated. These quiescent and slowly proliferating cells could be maintained in PDGFC supplemented media for up to 100 days. Furthermore, their behavior *in vitro* was stem-like, which is to say, they were able to self-renew in small minimally enlarging spheres, while also retaining the capacity to proliferate rapidly and/or differentiate in response to specific environmental cues. When switched from media supplemented with PDGFC to media containing EGF/FGF, they divided rapidly and continuously and expressed SOX2 and Nestin, and when FGF was added to PDGFC supplemented media they transitioned to cells expressing oligodendrocyte lineage markers.

Our assertion that sphere forming cells isolated from the SVZ using PDGFC are neural stem cells is also supported by their expression of stem cell markers, ability to express markers of neuronal, astrocytic and oligodendroglial differentiation when exposed to FBS, and analyses of bulk-RNA sequencing. The latter revealed that cells isolated and maintained in PDGFC, like those isolated in EGF/FGF, exhibit a stemness profile. Our other assertion that cultures isolated using PDGFC contain a mixture of quiescent and slowly proliferating cells, and hence are stem-like, is supported by immunofluorescence, cell cycle, and flow cytometry data in which we show that cells cultured in PDGFC are either non-dividing or dividing slowly. The quiescent nature of NSCs in PDGFC is reminiscent of type B cells of the SVZ (Doetsch et al., 1999) and their expression of the intermediate filament protein Nestin suggests a similarity to activated type B cells (Doetsch et al., 1999; Seroogy et al., 1995). This assumption is further supported by deconvolution analysis of our transcriptomic data with single-cell RNA-sequencing of the adult mouse SVZ (Cebrian-Silla et al., 2021) showing a significantly higher proportion of type B cells in the PDGFC condition *vs*. EGF/FGF and SVZ tissue. The mechanistic basis of the different proliferative behaviours of neural stem cells in PDGFC *vs*. EGF/FGF is associated with differences in their levels of expression of Cyclin D1 and D2 and the early response genes, *C-Jun* and *C-Fos*, all of which are under expressed in the PDGFC condition.

The PDGF ligands are among the most investigated growth factors owing to their interesting biology and versatile roles (Andrae et al., 2008). PDGFA has been extensively studied in the central nervous system and more recently PDGFC has been found to play a key role in both cardiovascular and neurobiology. While the expression of PDGFC and its receptor PDGFRα have been demonstrated in the adult brain, the effects of PDGFC on the regulation and proliferation of neural stem and progenitor cells has not been fully explored (Stefanitsch et al., 2015; Tian et al., 2021). Our findings suggest that PDGFC may play a role in regulating the composition of the murine SVZ, the major stem cell niche in the mammalian brain. Supporting this possibility is the observation that *Pdgfc* mRNA is highly expressed in adult murine SVZ tissue and is expressed at significantly higher levels in the SVZ than other regions of the mouse forebrain. In addition, PDGFC may contribute to the development of the SVZ. Ablation of PDGFC function has been demonstrated to affect the structural integrity of this region by causing ventricular malformations including asymmetry of the lateral ventricles, hypoplastic development of the septum separating the lateral ventricles, and loss of neuro-ependymal integrity, the epithelium from which NSCs emerge (Fredriksson et al., 2012; Stefanitsch et al., 2015)

In related work, we also observed that PDGFC in combination with FGF allows for the isolation, maintenance and propagation of OPCs from SVZ tissue and can replace PDGFA in this regard. This interesting result was reinforced by our observation that OPCs isolated traditionally in PDGFA/FGF and those isolated in PDGFC/FGF have indistinguishable gene expression profiles. Since NSCs do not survive in defined media supplemented with PDGFA alone, but do survive and proliferate in PDGFC, perhaps PDGFC is the more natural ligand for OPCs.

*In vitro* models using murine cells have limitations but many biological processes that proved to be relevant to human health were first observed in laboratory settings using simple systems. With that in mind, our work describes a method of culturing NSCs from the SVZ that relies on PDGFC. In PDGFC, pluripotent neural stem cells are quiescent and slowly proliferating, mimicking the behavior of Type B cells in the SVZ. This new laboratory model may facilitate experiments that probe a broad range of neural stem cell behaviours and their contribution to diseases of the brain, including cancers.

## Materials and Methods

### Mice and cell culture

NSC cultures were established from the SVZ of adult C57BL/6J mice as previously described (Reynolds and Weiss, 1992). Cultures were supplemented with PDGF-CC; Heparin; Epidermal Growth Factor; Fibroblast Growth Factor; or PDGF-AA. Primary cultures were cleared of debris by centrifugation and passaged when sphere diameter reached 100-200µm. Differentiation was induced by adhering spheres to poly-D-lysine and Laminin coated coverslips and incubation with 1% FBS for 6 days [see supplemental methods]. All experiments involving mice followed policies and procedures set by the Animal Care Committee at the University of Calgary (protocol #AC24-0094).

### Sphere diameter measurements

Images were taken using an EVOS cell imaging system. Image analysis and diameter measurements were performed using FIJI/ImageJ.

### EdU incorporation

Uptake was assessed using the Click-iT EdU Proliferation kit. Samples were incubated in 5-ethynyl-2′-deoxyuridine for 24hrs and stained as instructed by the manufacturer. Images were acquired on a ZEISS LSM880 microscope.

### Immunofluorescence

Cells were stained as previously described (Omairi et al., 2023). Cells were fixed using a PEM buffer, PFA and Glutaraldehyde and stained using the antibodies listed in supplemental methods. Images were acquired on a ZEISS AxioObserver or LSM880 microscopes. Image analysis and quantification used FIJI/ImageJ.

### Cell cycle analysis

NSCs were fixed with 95% ethanol, treated with ribonuclease then stained with Propidium Iodide. Cells were analyzed using a BD FACS Canto Flow Cytometer by PI pulse (Ex 493nm, Em 636nm). To fit Gaussian curves, ModFit LT V3.3.11 was used. Experiments were performed in triplicate with three independent replicates.

### Cell proliferation tracking with CFDA-SE

The rate of cell division was tracked using CFDA-SE [5(6)-carboxyfluorescein diacetate succinimidyl ester]. Cells were expanded in EGF/FGF and incubated with CFDA-SE solution, washed then excess stain was quenched in 10% FBS. Cells were cultured in EGF/FGF or PDGFC for 24, 48, 72, and 96hrs (n=5). Fluorescence was quantified using a BD FACS Canto Flow Cytometer (Ex 492nm, Em 517nm).

### Transcriptome sequencing

RNA was isolated from cells cultured in PDGFC (n=3), EGF/FGF (n=3), PDGFA+FGF (n=3) and PDGFC+FGF (n=3) for 7 days, or from fresh SVZ tissue (n=3) using the Qiagen RNeasy kit. RNA was assessed for quality using a Nanodrop™ and sent to the Centre for Health Genomics & Informatics at the University of Calgary for library preparation and sequencing (NovaSeq 6000; 50M reads/sample: rRNA depleted). Reads were aligned to the mm10 reference genome (GRCm38; Genome Reference Consortium) and counted (STARv.2.7.0a) [see supplemental information]. Gene expression differences, PCA plots, and heatmaps were generated using R-package DESeq2 v.1.26.0.

CIBERSORTx (Newman et al., 2019) was used to perform deconvolution of bulk RNA sequencing. The single cell reference matrix was generated from previously published single cell RNA sequencing of the adult mouse ventricular-subventricular zone (Cebrian-Silla et al., 2021) using the described cell types. The single cell RNA sequencing dataset was subset to 10000 cells due to size constraints in the CIBERSORTx web portal.

## Statistical analysis

Statistics were performed on GraphPad Prism version 8 or R studio version 3.6.3. One-way ANOVA with Tukey post-analysis for multiple comparisons within a sample set and unpaired t-test in sample sets of two groups were used. Two proportions z-test was used to compare two proportions. All tests were two-sided. Error bars are represented on graphs.

## Resource availability

### Lead contact

Requests for information may be directed to Dr. Cairncross.

### Materials availability

This study did not generate new unique reagents.

### Data availability

Transcriptomic (RNA-seq) data have been deposited at GEO at GEO: GSE288038 and will be publicly available as of the date of publication.

## Acknowledgments

HKO received graduate student awards from the Arnie Charbonneau Cancer Institute at the University of Calgary; SS received undergraduate student research awards from Alberta Innovates and the University of Calgary; JAC and JGC were supported by the Alberta Children’s Hospital Foundation Cancer Research Chair and the Brain Tumour Research Chair at the University of Calgary, respectively.

## Author contributions

HKO and SB made the observations upon which this manuscript is based; HKO and SS characterized the NSC and OPC cultures and performed the immunofluorescence experiments; MM and KMH performed the transcriptomic, bioinformatic, and cell cycle analyses. HKO, MM, SS, JAC, and JGC designed the experiments, interpreted the results, and wrote the manuscript.

## Declaration of interests

The authors declare no competing interests.

## Notes

### Competing Interest Statement

The authors have declared no competing interest.

