## Supplemental Information for "Isolation of Neural Stem Cells Using Platelet Derived Growth Factor C"

### Supplemental figures

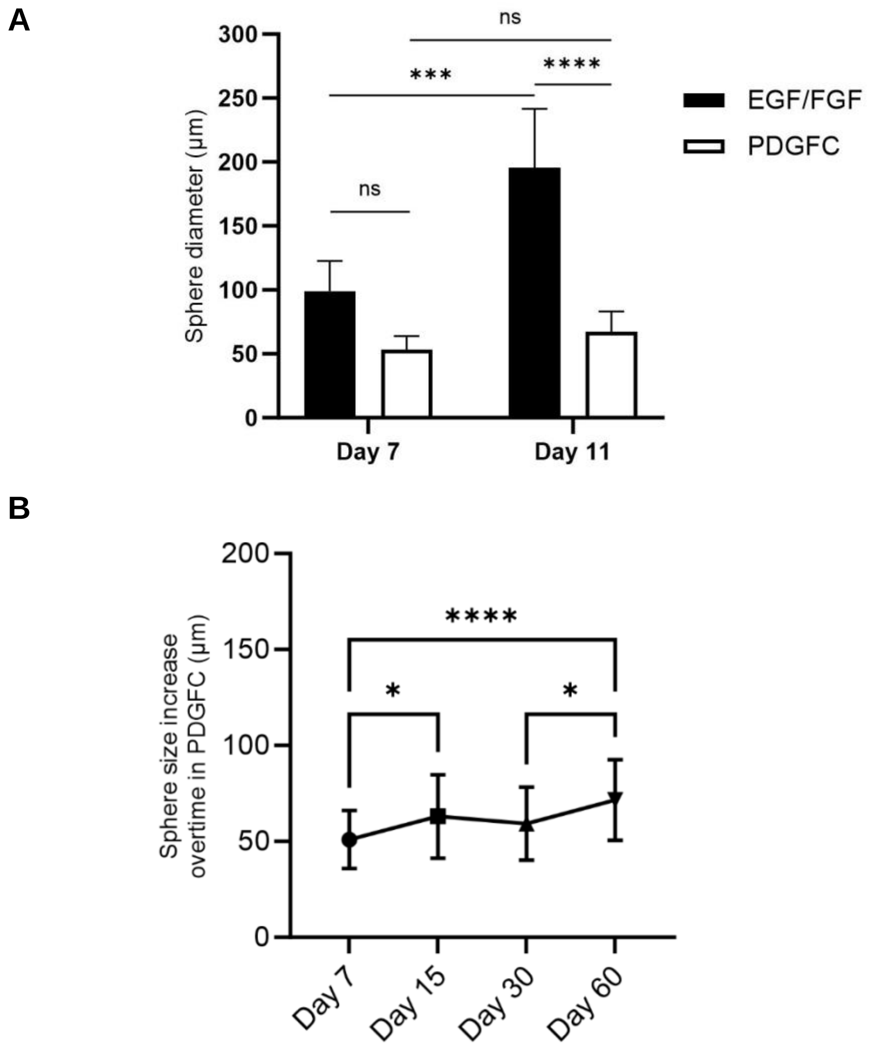


**Figure S1. SVZ-derived neural stem cells isolated using PDGFC**

(**A)** Bar graph showing sphere diameter in PDGFC and EGF/FGF cultures after 7 and 11 days. At day 7, 132 spheres were measured in PDGFC (n=4) and 96 spheres were measured in EGF/FGF (n=5); at day 11, 193 were measured in PDGFC (n=5) and 192 in EGF/FGF (n=6). **(B)** Histogram representing the diameter of spheres cultured in PDGFC at 5, 15, 30 and 60 days after the cultures were initiated. (ns= not significant; *= p <0.05; ****= p <0.0001)

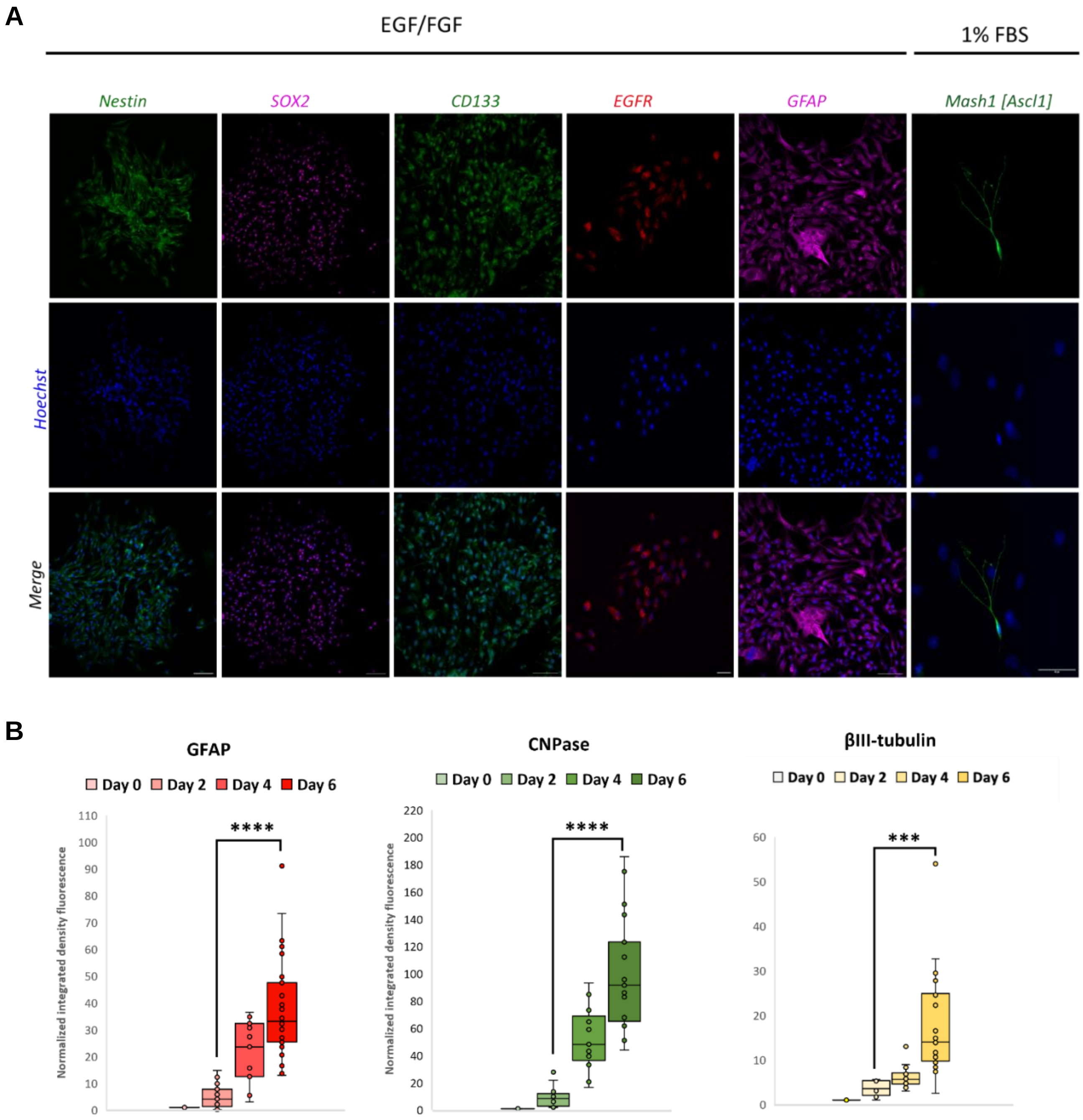


**Figure S2. Stem and differentiation markers in SVZ-derived neural stem cells isolated using EGF/FGF and PDGFC**

**(A)** Representative immunofluorescence images of SVZ-derived cells in EGF/FGF stained for the neural stem cell and type B markers Nestin, SOX2, CD133, EGFR and GFAP. Positive control for Type C marker Ascl1 [Mash1] using cells differentiated in 1% FBS for 4 days [Scale bar: 50 μm]. **(B)** Box blots showing increasing levels of GFAP, CNPase, and α-II-tubulin in NPCs in PDGFA during the process of differentiation in FBS. Statistical analyses were performed using paired t-test. (*** = p <0.001; ****= p <0.0001).

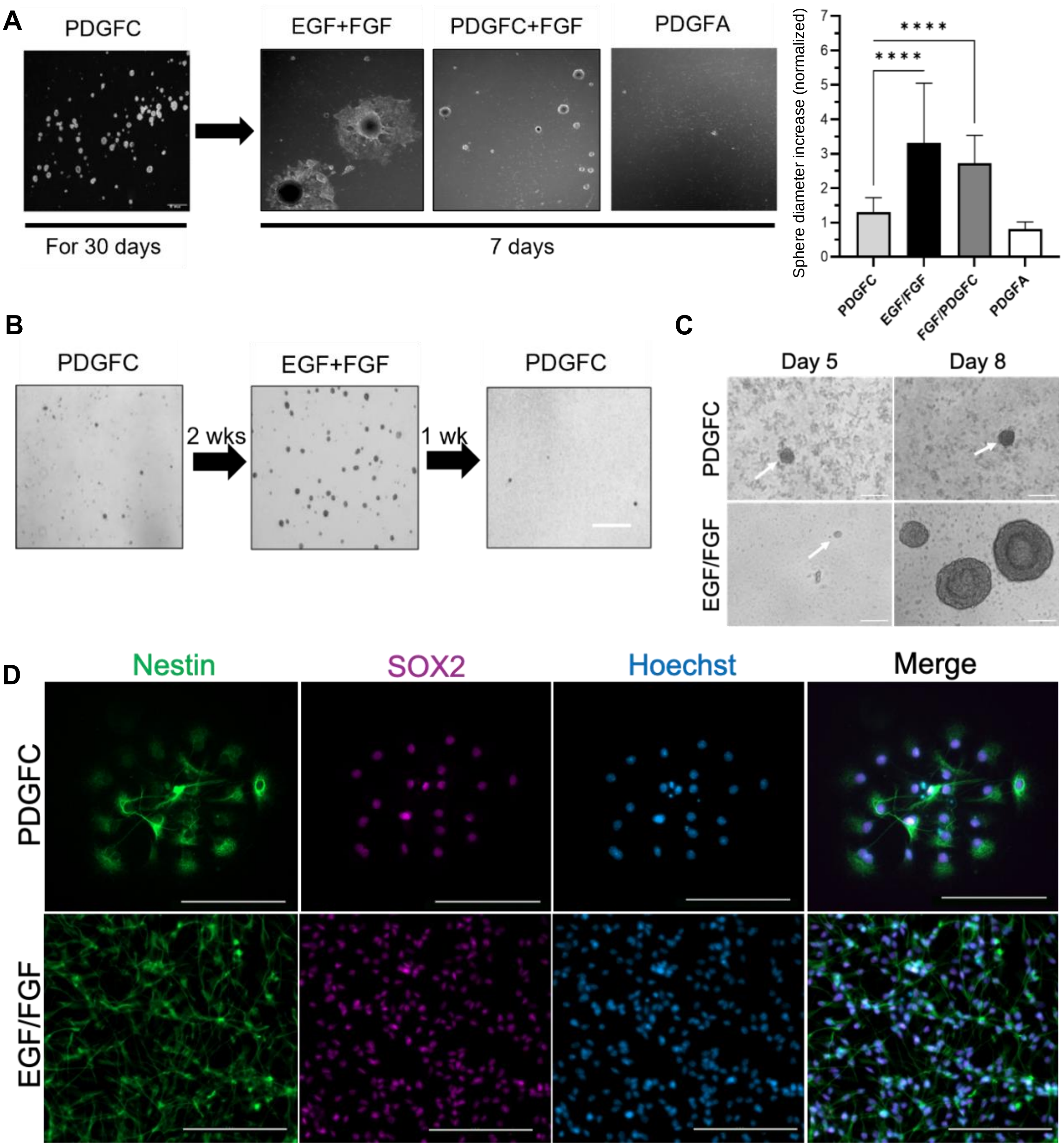


**Figure S3. NSCs in PDGFC can be induced to proliferate and can be isolated from the cortex**

(**A**) Bright field images of spheres isolated in PDGFC and switched to EGF/FGF, PDGFC+FGF, and PDGFA with measurement of sphere size (fold change, ****=p≤0.0001). **(B)** Bright field images of spheres in PDGFC switched to EGF/FGF for 2 weeks and switched back to PDGFC for 1 week. **(C)** Representative brightfield images of spheres in PDGFC or EGF/FGF isolated from the cortex **(D)** Representative immunofluorescence images of cortex-derived cells in PDGFC and EGF/FGF stained for neural stem cell markers Nestin, SOX2. [Scale bar: 100 µm].

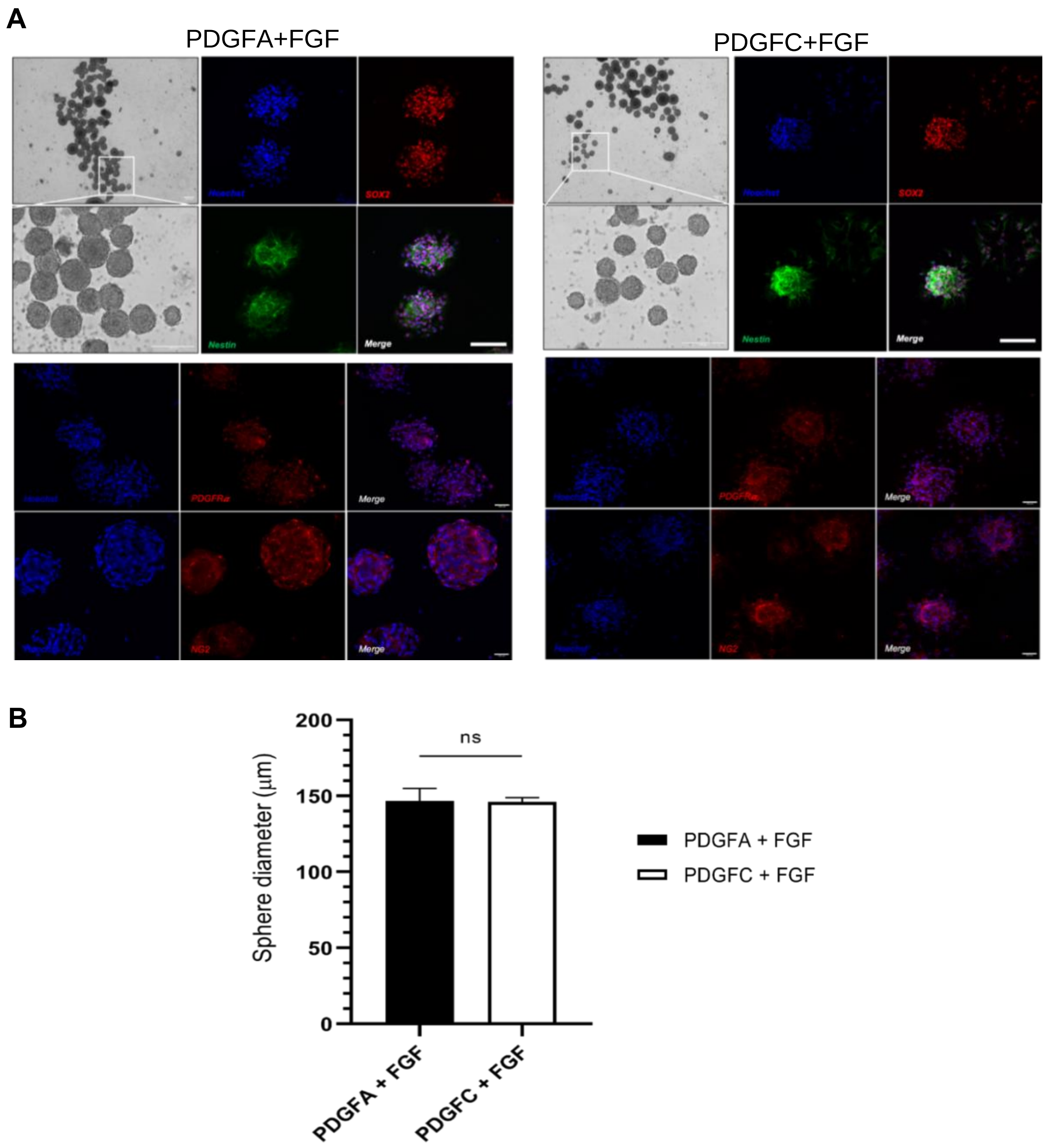


**Figure S4. PDGFC replaces PDGFA in OPC isolation protocols.**

**(A)** Representative bright field images and immunofluorescence images of SVZ-derived cells in PDGFC+FGF and PDGFA+FGF (control) stained for the stemness markers (SOX2 and Nestin), the glial marker (GFAP) and for markers of OPCs (PDGFRα and NG2). [Scale bar: 100 µm]. **(B)** Bar graph showing sphere diameter in PDGFA +FGF (n=3) and PDGFC+FGF (n=3) from primary cultures 7 days after SVZ microdissection (unpaired t-test, ns=not significant).

#### Supplemental methods

***Mice and cell culture***: C57BL/6J (stock # 000664) mice were purchased from The Jackson Laboratory. Cultures of NSCs were established from the SVZ of 8 to 12-week-old male and female mice as previously described (Azari et al., 2010; Reynolds and Weiss, 1992). Cells were supplemented with different growth factors including PDGF-CC (20ng/µL); Heparin (20ng/µl); Epidermal Growth Factor (20ng/µl); Fibroblast Growth Factor (20ng/µl); or PDGF-AA (20ng/µl). Primary cultures of SVZ-derived neurospheres were cleared of debris by centrifugation and passaged when the sphere diameter reached 100-200 µm. Primary NSC cultures were named as such: number of mouse-growth factor of culture-passage number (e.g. 27PDGFC-P4). No established/permanent cell lines were used. For passaging, cells were dissociated into single cells using Accumax and resuspended in media at a 1/10 ratio. To induce differentiation, spheres were adhered to glass cover slips coated with poly-D-lysine (PDL) (0.1 mg/ml) and Laminin (10 µg/ml) and incubated in 1% FBS for 6 days. All studies using mice were conducted in accordance with the policies and procedures established by the Animal Care Committee at the University of Calgary (protocol #M08029). In preparation for certain experiments, cells were maintained in media supplemented with EGF/FGF and then switched to PDGFC (i.e., experimental condition) or maintained in EGF/FGF (i.e., control condition). EGF/FGF cell lines were passaged under 7 times and discarded thereafter. For thawing, cells were cleaned of Dimethyl sulfoxide (DMSO) by centrifugation and resuspended in warmed intended media. Primary cell lines were not systematically tested for Mycoplasma.

***Sphere diameter measurements***: Bright field images of spheres were taken using an EVOS cell imaging system (10x magnification). Image analysis and diameter measurements were performed using FIJI/ImageJ. At day 7, 132 (n=4) and 96 (n=5) spheres were measure in PDGFC and EGF/FGF, respectively and at day 11, 193 (n=5) and 192 (n=6) spheres were measured in PDGFC and EGF/FGF, respectively. Statistical analysis was performed using ordinary one-way ANOVA and Tukey's multiple comparisons test. In cultures that were switched to PDGFC from EGF/FGF, 105 spheres were measured. Statistical analysis was performed using ordinary one-way ANOVA test.

***EdU incorporation***: Proliferation was assessed using the Click-iT EdU Proliferation kit (C10337). Samples were adhered on PDL/Laminin-coated sterile coverslips and maintained in PDGFC or EGF/FGF. Samples were incubated in 5-ethynyl-2′-deoxyuridine (EdU) for 24 hours and prepared for staining following the manufacturer's protocol. Images were acquired on a ZEISS LSM880 confocal microscope with a 20x/0.8 PlanApo objective

***Immunofluorescent staining***: Cells were fixed with 3.2% PFA w/v, 0.25% Glutaraldehyde (GA) in a PEM buffer (0.1M PIPES, 0.001M MgCl2x 6 H2O, 0.001M EDTA, 0.5% v/v Triton-X-100) for 10 minutes at room temperature (RT), rendered permeable in 0.5% v/v Triton X-100/1XPBS for 4 minutes, and incubated in a blocking solution (2% v/v Triton X-100, 3% w/v Bovine Serum Albumin (BSA), 5% v/v Goat serum and 0.2% v/v sodium azide-20% in 1X 1XPBS) for 1 hour at RT. Immunofluorescent staining was performed using the following primary antibodies diluted in blocking buffer and incubated over night at 4°C:: anti-EGFR (1/100); anti-CD133 (1/100); anti–sex determining region Y–box 2 (Sox2) (1/500); anti-Nestin (1/50); anti-NG2 (1/400); anti-PDGFRα (1/1000); anti-GFAP (1/400); anti-CNPase (1/1000); anti–β-III-tubulin

(1/100); anti-Ascl1(1/100); anti-CDKN1B/Kip1 p27 (1/50); and anti-Ki-67 (1/500). The following secondary antibodies were used: anti-rabbit IgG-Alexa Fluor 594 (1/1000); anti-mouse IgG-Alexa Fluor 488 (1/1000) together with the nuclear counterstain, Hoechst 33342. Coverslips were washed and mounted on glass slides using Fluormount-G mounting medium (Southern Biotechnology). Images were acquired on a ZEISS LSM880 confocal microscope with a 20x/0.8 PlanApo objective. Image analysis and protein quantification during differentiation was performed using FIJI/Image J. The level of GFAP, CNPase and β-III-tubulin were quantified in 150, 69 and 87 cells, respectively. Statistical analysis was performed using the paired t-test.

***Cell cycle analysis***: Our protocol was adapted from the Abcam flow cytometry cell cycle analysis with Propidium Iodide DNA staining (www.abcam.com/protocols/flow-cytometric-analysis-of-cell-cycle-with-propidium-iodide-dna-staining). NSCs were dispersed into single cells, re-suspended in media, and fixed with 95% Ethanol added to 1XPBS (1:1) drop by drop while vortexing. Samples were washed with 1XPBS and treated with ribonuclease (50µl of 100µg/ml) for 15 minutes. PI (200µl) was added (50µg/ml). Cells were analyzed using a BD FACS Canto Flow Cytometer in the University of Calgary Flow Cytometry Core Facility by PI pulse (Ex 493nm, Em 636nm). To fit Gaussian curves to each phase, ModFit LT V3.3.11 was used. Experiments were done in triplicate with three independent replicates.

***Cell proliferation tracking with CFDA-SE***: The rate of cell division was tracked using CFDA-SE (5(6)-carboxyfluorescein diacetate succinimidyl ester) (Stem cell technologies, Cat # 75003). Cells ($1X10^8$) were expanded in EGF/FGF, harvested, washed once with 1XPBS, and incubated with CFDA-SE solution (10uM/mL) for 5-10 minutes at 37°C. Uninternalized stain was quenched in 10% v/v FBS for 5 mins at 37°C. Cells were then centrifuged and the supernatant discarded. Stained cells were resuspended in media containing EGF/FGF (20 ng/ml) or PDGFC (20 ng/ml) for 24hrs, 48hrs and 72hrs in 5 cultures, and 96hrs in 2 of them. All staining was done in the dark. Cells were analyzed using a BD FACS Canto Flow Cytometer in the University of Calgary Flow Cytometry Core Facility using the FITC signal detector (Ex = 488 nm: Em = 530 nm). To fit Gaussian curves to each phase, ModFit LT V3.3.11 was used. Statistical analysis was performed using the unpaired t-test.

***Transcriptome sequencing:*** RNA was isolated from cells cultured in PDGFC (n=3), EGF/FGF (n=3), PDGFA+FGF (n=3) and PDGFC+FGF (n=3) for 7 days, or from fresh SVZ tissue (n=3) using the Qiagen RNeasy kit. RNA was assessed for quality using a Nanodrop™ and sent to the Centre for Health Genomics & Informatics at the University of Calgary for library preparation and sequencing (NovaSeq 6000; 50M reads/sample: rRNA depleted). Reads were aligned to the mm10 reference genome (GRCm38; Genome Reference Consortium) and counted (STARv.2.7.0a)(Dobin et al., 2013). Differential gene expression, PCA plots, and heatmaps were generated using R-package DESeq2 v.1.26.0. (Love et al., 2014).

CIBERSORTx (Newman et al., 2019) was used to perform deconvolution of bulk RNA sequencing. The single cell reference matrix was generated from previously published single cell RNA sequencing of the adult mouse ventricular-subventricular zone (Cebrian-Silla et al., 2021) using the described cell types. The single cell RNA sequencing dataset was subset to 10000 cells due to size constraints in the CIBERSORTx web portal.

***Statistical analysis***: Statistics were performed on Graphpad Prism version 8 or R studio version 3.6.3. One-way ANOVA with Tukey post-analysis for multiple comparisons within a sample set and unpaired t-test in sample sets of two groups were used. Two proportions z-test was used to compare two observed proportions. All tests were two-sided. Error bars are represented on graphs.

| REAGENT | SOURCE | IDENTIFIER | CONCENTRATION |
|---|---|---|---|
| **Chemicals and Growth factors** | | | |
| Accumax | Fisher Scientific | 509271 | |
| CFDA-SE | Stem cell technologies | Cat # 75003 | 10µM |
| Click-iT EdU | Thermofisher Scientific | C10337 | 10µM [EdU] |
| Epidermal Growth Factor | Stem Cell Technologies | 78006 | 20ng/ µL |
| Fibroblast Growth Factor | Stem Cell Technologies | 78003 | 20ng/ µL |
| Heparin | Stem Cell Technologies | 07980 | 20ng/ µL |
| Laminin | Sigma | L2020 | 10µg/mL |
| PDGF-AA | Stem Cell Technologies | 78095 | 20ng/µL |
| PDGF-CC | Stem Cell Technologies | 78168 | 20ng/ µL |
| Poly-D-lysine | Gibco | A39804-01 | 0.1mg/mL |
| Propidium Iodide | Sigma | P4170 | 50µg/mL |
| Ribonuclease | Sigma | | 100µg/mL |
| **Antibodies** | | | **DILUTION** |
| Ascl1 | Thermofisher Scientific | 14-5794-82 | 1/100 |
| CD33 | Abcam | ab19898 | 1/100 |
| CDKN1B/Kip1 p27 | Santa Cruse technologies | Sc-1641 | 1/50 |
| CNPase | Millipore-Sigma | MAB326 | 1/1000 |
| EGFR | Cell Signaling Technology | #4267 | 1/100 |
| GFAP | Stem cell technologies | 60128 | 1/400 |
| Ki-67 | Abcam | ab15580 | 1/500 |
| Mouse IgG-AF 488 | Thermofisher Scientific | A11001 | 1/1000 |
| Nestin | Santa Cruse technologies | Sc-23927 | 1/50 |
| NG2 | Millipore-Sigma | AB5320 | 1/400 |
| PDGFR-alpha | Cell Signaling Technology | #3164S | 1/1000 |
| rabbit IgG-AF 594 | Thermofisher Scientific | A11037 | 1/1000 |
| SOX2 | Cell Signaling Technology | #3728 | 1/500 |
| β-III-tubulin | Millipore-Sigma | MAB1637 | 1/100 |
| **Other** | | | |
| NEBNext® Ultra™ | New England Biolabs | #E7770 | N/A |

**Table S1.** List of reagents and antibodies
